# Aging reshapes cardiac extravascular leukocyte transcription through bidirectional remodeling during LPS-induced endotoxemia

**DOI:** 10.64898/2026.11.02.763785

**Authors:** Qing Yu, Hongyu Gao, Fang Fang, Jiayue Du, Meijing Wang

**Affiliations:** Department of Surgery, Center for Surgical Sciences, Indiana University School of Medicine, Indianapolis, IN, USA; Center for Medical Genomics, Indiana University School of Medicine, Indianapolis, IN, USA

**Keywords:** Aging, Sepsis, Cardiac inflammation, Leukocytes, Macrophages, Lymphocytes, single-cell RNA sequencing (scRNA seq), LPS

## Abstract

Sepsis disproportionately affects older adults and is frequently complicated by cardiac dysfunction. However, how aging alters the transcriptional responses of cardiac extravascular immune cells during systemic inflammation remains poorly understood. Here, we investigated aging-dependent transcriptional remodeling of cardiac immune cells using an LPS-induced endotoxemia model in young and aged C57BL/6 mice. Mice received intraperitoneal LPS (5 mg/kg body weight) or vehicle, and cardiac extravascular leukocytes were isolated from ex vivo retrogradely perfused hearts 24 h later for single-cell RNA sequencing. Aging was associated with bidirectional transcriptional remodeling, characterized by both up- and downregulation of gene programs, particularly in cardiac macrophages, monocyte-like macrophages, T cells, and B cells following LPS challenge. A shared feature across these four cell populations was reduced representation of mitochondrial-associated transcriptional programs, indicating aging-dependent remodeling of mitochondrial biology across cardiac myeloid and lymphoid lineages during systemic inflammation. In parallel, cell type-specific transcriptional remodeling revealed distinct functional-associated responses. Macrophages and monocyte-like macrophages exhibited increased tissue-remodeling and repair-associated transcriptional programs. Both T and B cells showed reduced metabolic programs but increased actin cytoskeleton organization. T cells additionally exhibited reduced cytotoxicity-associated and enhanced Th17-associated transcriptional programs, whereas B cells showed reduced antigen-processing and presentation programs. Cell-cell communication analysis suggested aging-associated differences in immune cell interactions following LPS challenge and potential features of cardiac inflammaging at baseline. Together, these findings demonstrate that aging bidirectionally reshapes the cardiac immune response to systemic inflammation, characterized by shared remodeling of mitochondrial-associated programs and cell type-specific changes in inflammatory, metabolic, cytoskeletal, and effector-associated programs. These aging-dependent transcriptional changes occurred in the context of the greater severity of illness observed in aged mice following LPS-induced endotoxemia.

## INTRODUCTION

Sepsis is a life-threatening syndrome characterized by a dysregulated host response to infection that can result in widespread tissue injury and multiple organ dysfunction [1–4]. Although substantial progress has been made in understanding the systemic inflammatory and circulatory responses associated with sepsis, increasing evidence indicates that immune responses within individual organs are important determinants of tissue injury, dysfunction, and repair [5]. Major organs possess distinct immune microenvironments that undergo organ-specific recruitment, activation, and functional reprogramming of resident and infiltrating immune cells during systemic inflammation. Thus, defining tissue-specific immune responses may provide important insight into the mechanisms underlying organ dysfunction during sepsis.

The heart is particularly vulnerable to systemic inflammatory injury. Septic cardiomyopathy, characterized by acute myocardial dysfunction that can be reversible but is associated with adverse clinical outcomes, occurs frequently in patients with severe sepsis or septic shock [6–9]. The cardiac immune-cell milieu, comprising resident and infiltrating leukocytes within the myocardial interstitium, may provide important clues to how the heart responds to systemic inflammation. Experimental models are instrumental in defining the cellular and molecular responses of the heart during systemic inflammatory challenges. The cardiac immune compartment comprises diverse populations of resident and recruited leukocytes, which collectively contribute to inflammatory signaling, cytokine production, tissue remodeling, and repair [10–13]. Recent single-cell studies have further revealed heterogeneous cardiac immune-cell states during experimental sepsis, including specialized macrophage populations associated with the septic cardiac microenvironment [14].

A major methodological challenge in characterizing the cardiac immune response during systemic inflammation is distinguishing leukocytes associated with cardiac tissue from those contained within the cardiac vasculature. Because the heart is highly vascularized, conventional tissue dissociation can recover both extravascular tissue-associated leukocytes and intravascular circulating cells, complicating the interpretation of local cardiac immune responses. Ex vivo retrograde perfusion provides an approach to substantially reduce the contribution of circulating blood cells before tissue collection, thereby enriching for leukocytes associated with the extravascular cardiac compartment. This distinction is particularly important during systemic inflammation, when the abundance, composition, and activation states of circulating leukocytes can change substantially. Characterizing the extravascular cardiac immune compartment can therefore provide a more tissue-proximal view of the cellular responses occurring within the heart during systemic inflammatory challenge.

Older adults bear a disproportionate burden of sepsis, with increased susceptibility to severe disease and substantially higher mortality than younger individuals [15–17]. Aging is also associated with an increased risk of organ dysfunction and adverse outcomes during sepsis [18–22], potentially reflecting aging-associated alterations in immune and tissue homeostasis. Inflammaging, broadly characterized by elevated basal inflammatory activity and progressive remodeling of innate and adaptive immune function, is a prominent feature of aging [23–25]. This state has been linked to the cumulative effects of cellular senescence, tissue damage, altered metabolic homeostasis, mitochondrial dysfunction, and persistent endogenous inflammatory stimuli [26]. At the same time, aging is associated with impaired immune responsiveness, altered leukocyte differentiation, and diminished capacity to appropriately resolve inflammatory responses [27, 28]. Consequently, the aged immune system may respond differently to acute systemic inflammatory challenges, with aging-associated alterations in the inflammatory baseline potentially influencing the magnitude, composition, and resolution of tissue-specific immune responses.

These aging-related changes may be particularly relevant in the heart, where older patients are more vulnerable to cardiac dysfunction during sepsis [29–32]. However, it is unclear how aging affects the response of extravascular cardiac leukocytes, which directly reflect the immune milieu within the cardiac tissue, during an acute systemic inflammatory challenge such as in sepsis. In the present study, we used an LPS-induced endotoxemia model to investigate how aging modifies the cardiac extravascular immune response to systemic inflammation. Cardiac extravascular leukocytes were isolated from ex vivo retrogradely perfused hearts of young and aged mice 24 h after LPS administration and profiled using single-cell RNA sequencing (scRNA-seq). To distinguish aging-dependent modification of the inflammatory response from baseline differences associated with aging or LPS exposure alone, we used a pseudobulk age × LPS interaction model to identify genes whose transcriptional responses to LPS differed between young and aged mice. Together, these approaches provide a single-cell view of the cardiac extravascular immune response and identify aging-dependent transcriptional programs associated with systemic inflammation.

## MATERIALS AND METHODS

### Animals and LPS administration

Male C57BL/6J mice at 9 weeks (young) or 90 weeks (aged) of age were obtained from the Jackson Laboratory and housed under specific pathogen-free (SPF) conditions at the Indiana University Laboratory Animal Resource Center (LARC). All animal procedures were performed in accordance with protocols approved by the Institutional Animal Care and Use Committee (IACUC). The mice received intraperitoneal injections of lipopolysaccharide (LPS; Sigma-Aldrich, L3129-100MG) at a dose of 5 mg/kg or an equivalent volume of phosphate-buffered saline (PBS) as control. Twenty-four hours after the injection, the mice were euthanized. Four experimental groups were analyzed: Young-control, Young-LPS, Aged-control, and Aged-LPS. Four biological replicates were initially generated for each experimental group. One Aged-LPS replicate was excluded from downstream analyses because the animal experienced cardiac arrest before euthanasia and required resuscitation, which was considered a potential confounding physiological perturbation affecting cardiac immune cell transcriptional states.

### Heart tissue collection and cardiac leukocyte isolation

The mice were euthanized and the hearts were harvested. The hearts were subjected to retrograde perfusion [33], minced, and enzymatically digested in digestion buffer containing collagenase type II (Worthington Biochemical, LS004176; 2 mg/ml) and Ca^++^ (0.5 mM) at 37°C for 1 hour. During digestion, tubes were inverted every 10 minutes to facilitate tissue dissociation. The resulting cell suspension was filtered through 70-μm cell strainer. The cells were washed and resuspended in MACS buffer (Miltenyi Biotec), followed by enrichment of leukocytes using mouse CD45 MicroBeads (Miltenyi Biotec, 130-052-301) according to the manufacturer’s protocol.

### Single-cell RNA sequencing library preparation & Sequencing

To enable multiplexing of biological samples, hashtag oligonucleotide (HTO)-conjugated antibodies were used for sample labeling. The following TotalSeq™-B anti-mouse hashtag antibodies (BioLegend) were used: TotalSeq™-B0301 anti-mouse Hashtag 1 Antibody, barcode sequence: ACCCACCAGTAAGAC; TotalSeq™-B0302 anti-mouse Hashtag 2 Antibody, barcode sequence: GGTCGAGAGCATTCA; TotalSeq™-B0304 anti-mouse Hashtag 4 Antibody, barcode sequence: AAAGCATTCTTCACG; TotalSeq™-B0306 anti-mouse Hashtag 6 Antibody, barcode sequence: TATGCTGCCACGGTA. Single-cell libraries were generated using the Chromium Next GEM Single Cell 3′ Kit v3.1 (10x Genomics, PN 1000269) and Chromium Next GEM Chip G Single Cell Kit (10x Genomics, PN 1000127) according to the manufacturer’s instructions. Single-cell RNA sequencing libraries were sequenced on an Illumina sequencer.

### Single-cell RNA sequencing preprocessing

Raw sequencing data were processed using Cell Ranger version 9.0.1 (10x Genomics). Reads were aligned to the mouse reference transcriptome (mm10-2020-A), and gene expression and feature-barcode matrices were generated using the Cell Ranger pipeline. Filtered feature-barcode matrices containing gene expression and HTO counts were imported into R (version 4.5.2) and analyzed using Seurat version 5.3.1.

### HTO demultiplexing & Quality control filtering

HTO count matrices were analyzed using the HTODemux workflow implemented in Seurat. HTO counts were normalized using centered log-ratio (CLR) normalization, followed by sample demultiplexing. Only cells classified as singlets were retained for downstream analyses. To prevent barcode duplication during integration of multiple sequencing pools, unique prefixes were added to cell barcodes from each pool prior to dataset merging. One biological replicate from Aged-LPS group was excluded following HTO demultiplexing, because the animal experienced cardiac arrest before euthanasia and required resuscitation, which was considered a potential confounding physiological perturbation affecting cardiac immune cell transcriptional states. Quality control filtering was performed to remove low-quality cells and potential technical artifacts.

### Data preprocessing, dimensional reduction, clustering, and cell type annotation

Gene expression matrices were normalized using the NormalizeData() function in Seurat. Highly variable genes were identified using FindVariableFeatures(), followed by data scaling using ScaleData(). Principal component analysis (PCA) was performed for dimensionality reduction, followed by construction of a shared nearest-neighbor graph using FindNeighbors(). Uniform Manifold Approximation and Projection (UMAP) was performed for visualization, and graph-based clustering was performed using FindClusters(). Cell clusters were annotated based on the expression of established canonical marker genes for immune cell populations. The annotated immune cell populations were subsequently used for differential expression and pathway analyses.

### Differential gene expression analysis

Differential gene expression analysis was performed using a pseudobulk approach, in which gene counts were aggregated by biological replicate within each annotated immune cell population. Pseudobulk count matrices were analyzed using DESeq2 with a design formula including age, treatment, and their interaction (∼ age + treatment + age:treatment). The age × treatment interaction term was used to identify genes exhibiting differential transcriptional responses to LPS between young and aged mice. Genes with an adjusted P value <0.05 and an absolute log2 fold change >1 were considered significant interaction differentially expressed genes. Volcano plots were generated using ggplot2. Up-INT genes were defined as genes with positive age × treatment interaction coefficients, indicating a more positive (or less negative) LPS-associated log2 fold change in aged mice compared with young mice. Conversely, Down-INT genes exhibited negative interaction coefficients, indicating a more negative (or less positive) LPS-associated log2 fold change in aged mice.

### Heatmap visualization

Heatmaps were generated to visualize significant age × LPS interaction differentially expressed genes within each immune cell population. Genes meeting the criteria of adjusted P value <0.05 and absolute log2 fold change >1 were ranked by adjusted P value, and the top 30 genes were selected for visualization. Heatmaps were generated using pheatmap package, with hierarchical clustering applied to genes and samples displayed in predefined experimental order (Young control, Young LPS, Aged control, and Aged LPS).

### Gene Ontology enrichment analysis

Gene Ontology (GO) Biological Process enrichment analysis was performed separately for upregulated and downregulated age × LPS interaction differentially expressed genes identified within each immune cell population. Enrichment analysis was performed using the clusterProfiler package (version 4.18.4) with the enrichGO() function. Analyses were restricted to the Biological Process ontology, and statistical significance was assessed using adjusted P values. Enriched pathways were visualized using dot plots generated with enrichplot (version 1.30.5), displaying the top 10 significantly enriched biological processes.

### Gene expression visualization

Expression of selected genes was visualized using violin plots generated with the VlnPlot() function in Seurat. Plots were generated separately for each annotated immune cell population and displayed single-cell expression patterns across the four experimental groups.

### Cardiac extravascular immune cell communication analysis using CellChat

Cell–cell communication among cardiac extravascular immune cell populations was inferred using CellChat. Seurat objects corresponding to the four experimental groups (Young-control, Young-LPS, Aged-control, and Aged-LPS) were analyzed separately using normalized gene expression data, cell-type annotations, and the mouse ligand–receptor interaction database provided by CellChat. Circle plots and heatmaps were generated to visualize the inferred communication networks and interaction strengths among immune cell populations in each group.

## RESULTS

### LPS and aging remodel the cardiac extravascular immune landscape

To characterize the cardiac extravascular immune landscape during endotoxemia, single-cell RNA sequencing (scRNA-seq) was performed on CD45-enriched cells isolated from retrogradely perfused hearts of young and aged mice 24 h after vehicle or LPS administration. The experimental design and analytical workflow are shown in Figure 1. Notably, following administration of LPS at 5 mg/kg, young mice remained relatively healthy and active throughout the 24-h period, whereas aged mice exhibited pronounced signs of illness prior to sacrifice. Cell identities were assigned based on the expression of canonical marker genes, revealing macrophages, monocyte-like macrophages, neutrophils, dendritic cells, NK cells, T cells, and B cells as the major immune populations (Figure 2A). Small populations of endothelial cells and fibroblasts were also detected, likely reflecting minor co-elution during CD45-based magnetic enrichment.

**Figure 1.**
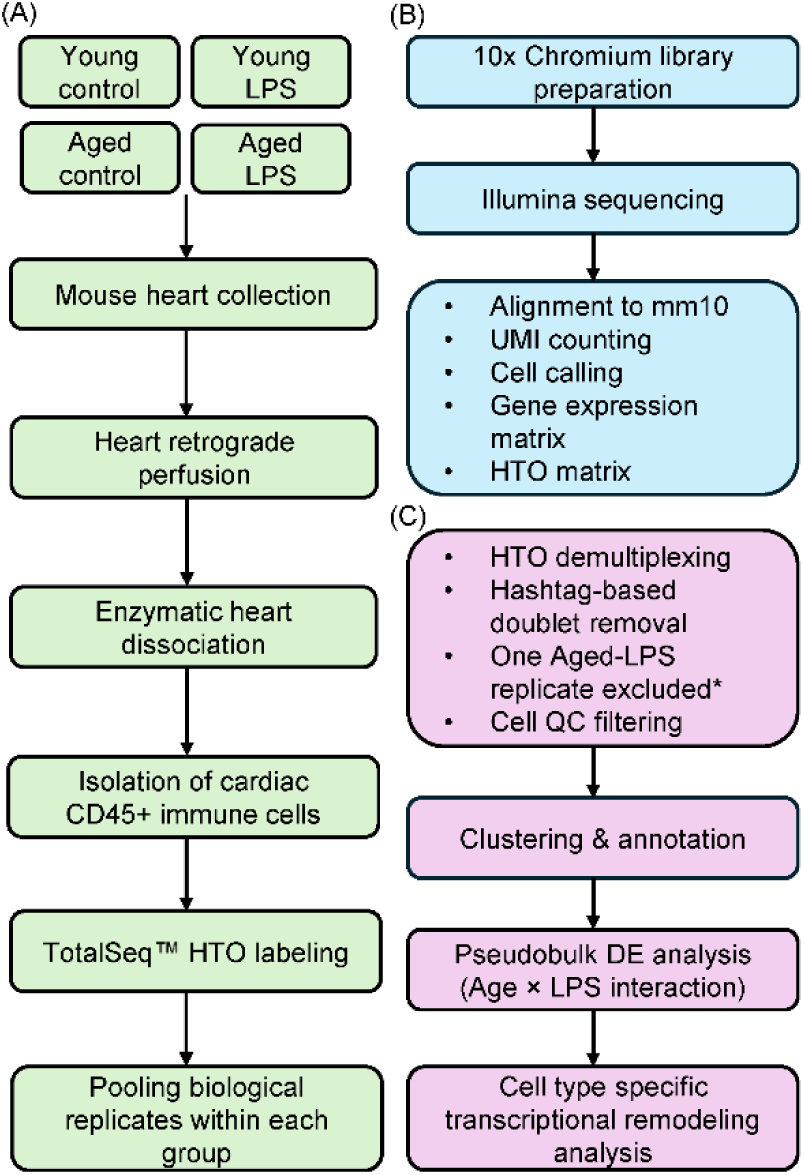
Experimental workflow and analytical pipeline. (A) Young (9-week-old) and aged (90-week-old) mice received vehicle control or lipopolysaccharide (LPS) 24 h before euthanasia. Hearts were collected, retrogradely perfused, and enzymatically dissociated to isolate extravascular CD45⁺ cardiac immune cells. Cells were labeled with TotalSeq™ hashtag oligonucleotide (HTO) antibodies and multiplexed for single-cell RNA sequencing (scRNA-seq), with n = 4 biological replicates per experimental group at the start of the study. (B) Single-cell libraries were generated using the 10x Genomics Chromium Single Cell 3′ Gene Expression platform, followed by Illumina sequencing and Cell Ranger processing to generate gene-expression and HTO feature-barcode matrices. (C) Downstream analyses included HTO demultiplexing, hashtag-based doublet removal, quality-control (QC) filtering, clustering, and cell-type annotation. Pseudobulk differential-expression analysis using the age × LPS interaction term was subsequently performed to identify cell type-specific aging-dependent transcriptional responses to LPS. *One Aged-LPS biological replicate was excluded from downstream analysis because the mouse experienced cardiac arrest before euthanasia, representing a potential biological confounder. The final sample size analyzed was therefore n = 4 for Young-control, Young-LPS, and Aged-control groups and n = 3 for the Aged-LPS group.

**Figure 2.**
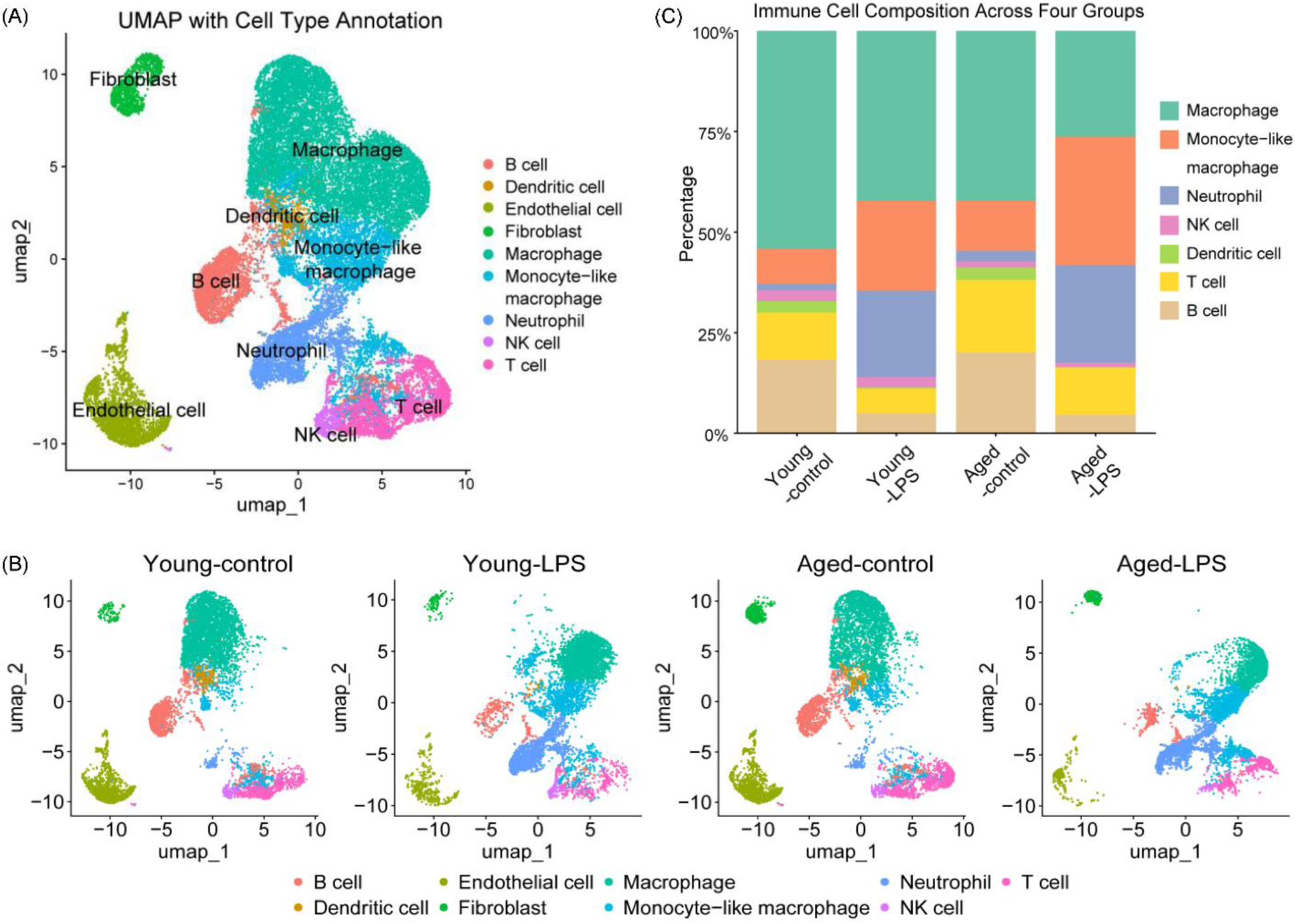
Clustering, annotation, and cellular composition of cardiac extravascular immune cells. (A) UMAP visualization of the integrated scRNA-seq dataset showing the major cardiac extravascular immune cell populations identified by clustering and canonical marker-gene expression. (B) UMAP visualizations showing the distribution of immune cell populations within each experimental group, illustrating group-dependent changes in the cardiac extravascular immune landscape. (C) Stacked bar plots showing the relative abundance of identified cell populations across the four experimental groups.

UMAP analysis showed that the cellular landscapes of young and aged control hearts were broadly similar, whereas LPS treatment substantially altered the cardiac immune landscape in both age groups (Figure 2B). The cellular composition of LPS-treated hearts also differed between young and aged mice, indicating that aging modified the cellular response to endotoxemia. Consistent with these global changes, LPS challenge altered the relative abundance of several immune populations (Figure 2C). The proportion of macrophages decreased following LPS treatment, with a greater reduction in aged hearts. In contrast, monocyte-like macrophages increased markedly after LPS administration, particularly in aged mice. Neutrophils were scarce in control hearts but increased substantially following LPS challenge. NK-cell proportions remained relatively stable and low across experimental groups, although they were slightly higher in young mice than in aged mice. Dendritic cells were nearly absent following LPS treatment, whereas the proportions of both T cells and B cells decreased following endotoxemia.

Together, these findings demonstrate that LPS exposure substantially remodeled the cellular composition of the cardiac immune landscape and that aging further modified this response. The scarcity of neutrophils in control hearts also supports effective removal of intravascular blood during cardiac perfusion.

### Aging differentially modifies LPS-responsive gene transcription across cardiac immune populations

To identify transcriptional responses to LPS that differed by age, differential expression analysis was performed using the age × treatment interaction term. This interaction identified genes for which the magnitude or direction of the LPS-induced transcriptional response differed between young and aged mice, while accounting for baseline differences associated with age.

The number of interaction-associated differentially expressed genes (interaction DEGs) varied substantially among immune populations (Figure 3A). Macrophages contained 1,140 interaction DEGs, followed by 1,061 in monocyte-like macrophages, 1,837 in T cells, and 762 in B cells. In comparison, neutrophils, NK cells, and dendritic cells showed 196, 336, and 60 interaction DEGs, respectively. Based on the extent of aging-dependent transcriptional remodeling and their relative abundance within the cardiac immune compartment, subsequent analyses focused on macrophages, monocyte-like macrophages, T cells, and B cells.

**Figure 3.**
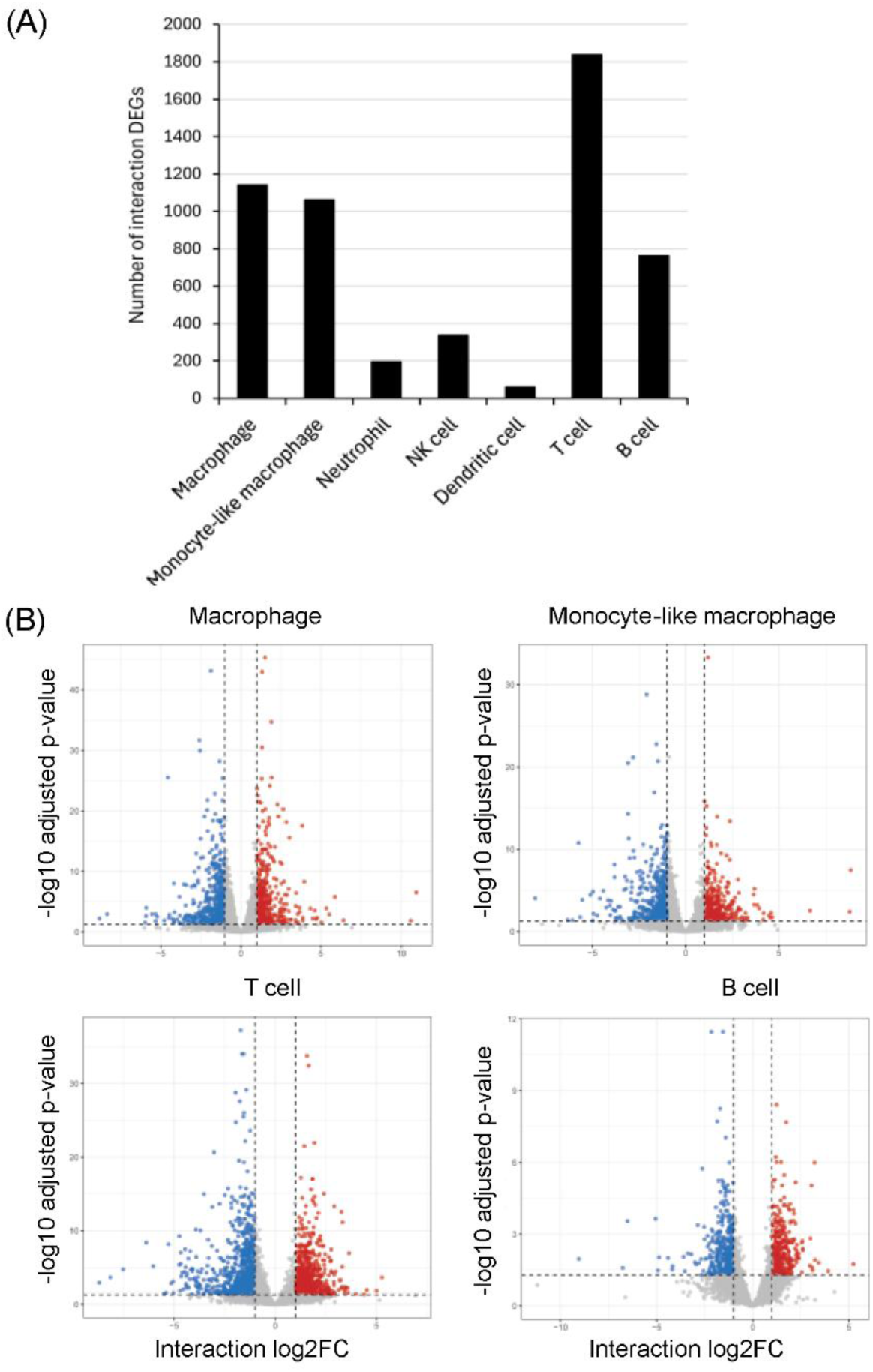
Age × LPS interaction differentially expressed genes across cardiac immune cell populations. (A) Bar plot showing the number of age × LPS interaction differentially expressed genes (interaction DEGs) identified in each cardiac immune cell population. (B) Volcano plots showing age × LPS interaction DEGs in macrophages, monocyte-like macrophages, T cells, and B cells. Red and blue dots indicate Up-INT and Down-INT genes, respectively. Up-INT genes were defined as genes with positive age × treatment interaction coefficients, indicating a more positive (or less negative) LPS-associated log2 fold change in aged mice compared with young mice. Conversely, Down-INT genes exhibited negative interaction coefficients, indicating a more negative (or less positive) LPS-associated log2 fold change in aged mice.

Volcano plots demonstrated both positive and negative interaction effects across all four focal populations (Figure 3B), with the magnitudes of positive and negative effects being relatively symmetrical within each cell population. These findings indicate that aging altered the LPS-responsive transcriptional programs in both directions rather than predominantly attenuating or compressing the transcriptional response.

### Aging remodels inflammatory and mitochondrial-associated programs in cardiac macrophages and monocyte-like macrophages

We examined aging-dependent modification of the LPS response in macrophages. The top 30 interaction DEGs are shown in Figure 4A. C1qb, a gene commonly associated with tissue-resident macrophages, showed an aging-dependent attenuation of its response to LPS. Spi1 (PU.1), a transcription factor involved in myeloid differentiation and macrophage lineage specification and maintenance, also exhibited a negative interaction effect. In contrast, Mrc1, which is associated with tissue-reparative macrophage states, showed a positive interaction effect. It is noteworthy that several mitochondrial DNA-encoded transcripts, including mt-Nd1, mt-Nd3, mt-Nd4, mt-Co2, mt-Co3, and mt-Atp6, showed positive age × LPS interaction effects. These genes encode components of mitochondrial respiratory complexes I, IV, and V. Their expression displayed a group-dependent pattern rather than a uniform increase across experimental groups, with particularly high expression in young-control and aged-LPS macrophages. GO enrichment analysis identified distinct functional categories among genes with positive and negative interaction effects (Figure 4B,C). Up-INT genes were enriched in pathways related to leukocyte chemotaxis, cell migration, oxidative stress responses, and JNK signaling. Down-INT genes were enriched in TNF production, myeloid leukocyte activation, inflammatory regulation, leukocyte proliferation, and antiviral responses.

**Figure 4.**
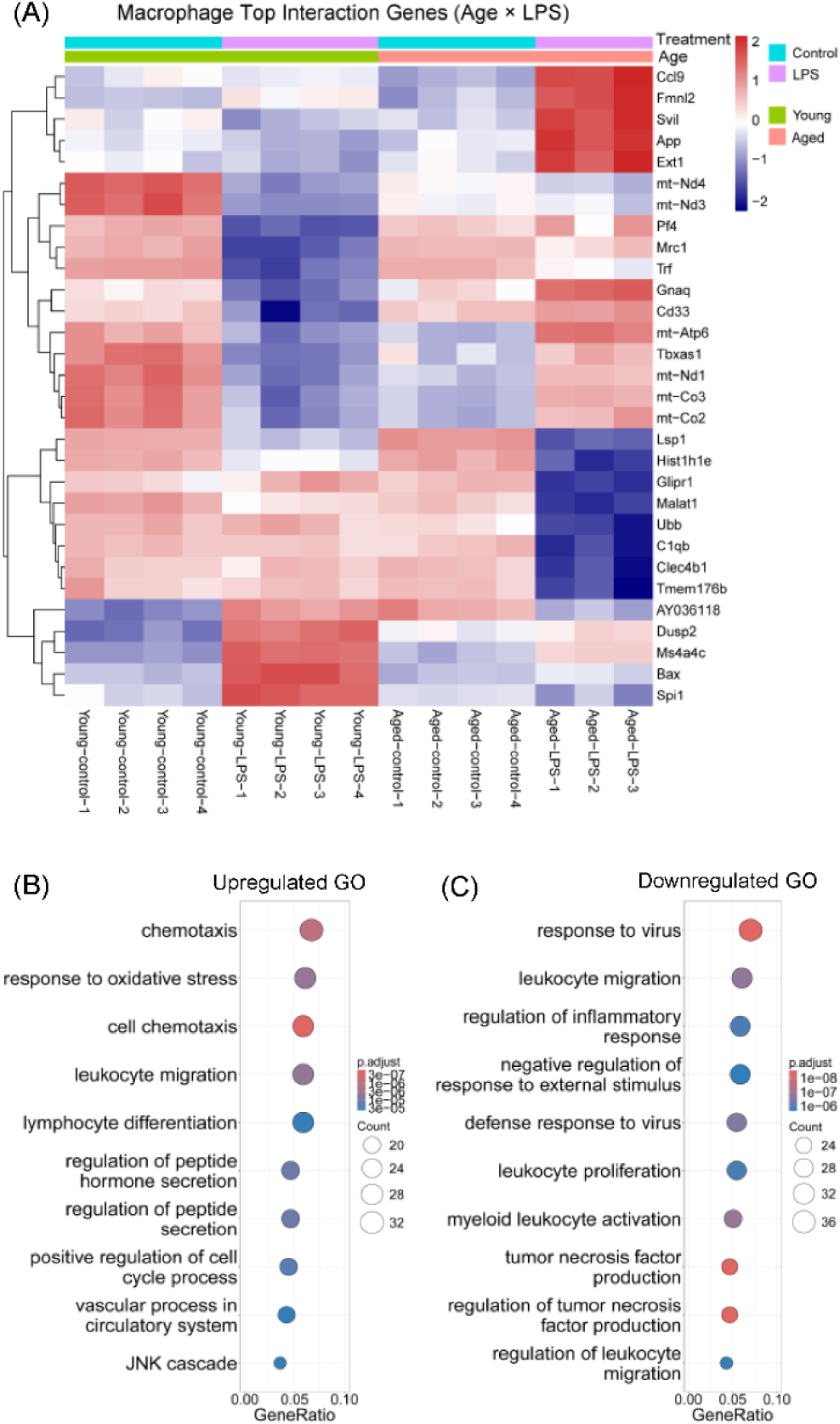
Age × LPS interaction differential expression and functional enrichment analyses in cardiac macrophages. (A) Pseudobulk heatmap showing the top 30 age × LPS interaction DEGs in macrophages. Genes meeting the criteria of adjusted P value < 0.05 and absolute log2 fold change > 1 were ranked by P value, and the top 30 genes were selected for visualization. (B, C) Gene Ontology (GO) enrichment analysis of Up-INT genes (B) and Down-INT genes (C). Up-INT genes were defined as genes with positive age × treatment interaction coefficients, indicating a more positive (or less negative) LPS-associated log2 fold change in aged mice compared with young mice. Conversely, Down-INT genes exhibited negative interaction coefficients, indicating a more negative (or less positive) LPS-associated log2 fold change in aged mice.

Monocyte-like macrophages likewise underwent extensive age-dependent transcriptional remodeling following LPS challenge (Figure 5A). Spi1 again exhibited a negative interaction effect, while several mitochondrial DNA-encoded genes showed positive interaction effects, consistent with the pattern observed in macrophages. GO enrichment analysis (Figure 5B,C) showed that Up-INT genes were enriched in T-cell activation, lymphocyte differentiation, cell adhesion, and hematopoietic regulation. In contrast, Down-INT genes were enriched in TNF production, nitric oxide biosynthesis, reactive oxygen species metabolism, myeloid leukocyte migration, mitochondrial organization, and apoptosis.

**Figure 5.**
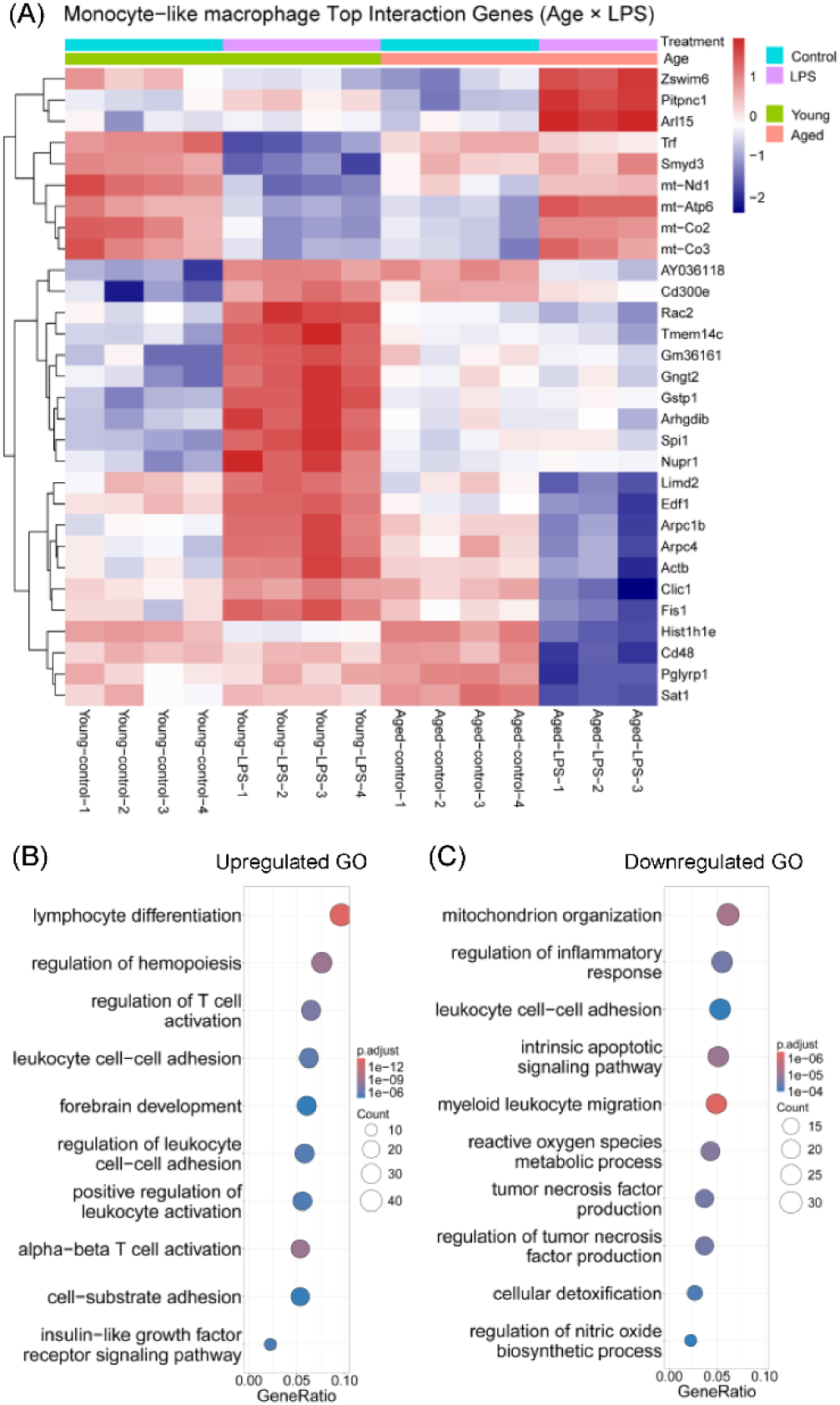
Age × LPS interaction differential expression and functional enrichment analyses in cardiac monocyte-like macrophages. (A) Pseudobulk heatmap showing the top 30 age × LPS interaction DEGs in monocyte-like macrophages. Genes meeting the criteria of adjusted P value < 0.05 and absolute log2 fold change > 1 were ranked by adjusted P value, and the top 30 genes were selected for visualization. (B, C) Gene Ontology (GO) enrichment analysis of Up-INT genes (B) and Down-INT genes (C).

To more comprehensively characterize these differences, we compared the full GO enrichment profiles of macrophages and monocyte-like macrophages and summarized the overlapping and distinct enriched GO terms in Table 1. Both populations showed enrichment of TNF production and type I interferon responses among Down-INT genes. Chemotaxis-related pathways were enriched among both Up-INT and Down-INT genes in both populations. Phagocytosis-related pathways were enriched among both Up-INT and Down-INT genes in macrophages but were predominantly associated with Down-INT genes in monocyte-like macrophages. Both populations also showed enrichment of tissue-remodeling and protein-phosphorylation pathways among Up-INT genes.

**Table 1.** Aging-dependent transcriptional remodeling of cardiac macrophages and monocyte-like macrophages during LPS-induced sepsis: GO enrichment of Up-INT DEGs and Down-INT DEGs.

| GO term | Macrophage |  |  | Monocyte-like macrophage |  |  |
|---|---|---|---|---|---|---|
|  | Down-INT | UP-INT | Summary | Down-INT | UP-INT | Summary |
| TNF production | ✓ | – | ↓ | ✓ | – | ↓ |
| Phagocytosis | ✓ | ✓ | ↓↑ | ✓ | – | ↓ |
| Response to virus | ✓ | ✓ | ↓↑ | ✓ | – | ↓ |
| Response to type I interferon | ✓ | – | ↓ | ✓ | – | ↓ |
| Response to type II interferon | ✓ | – | ↓ | ✓ | ✓ | ↓↑ |
| Chemotaxis | ✓ | ✓ | ↓↑ | ✓ | ✓ | ↓↑ |
| Tissue remodeling | – | ✓ | ↑ | – | ✓ | ↑ |
| Protein phosphorylation | – | ✓ | ↑ | – | ✓ | ↑ |
| Regulation of hemopoiesis | – | – | – | – | ✓ | ↑ |
| Oxidative phosphorylation | – | – | – | ✓ | – | ↓ |
| Mitochondrion organization | ✓ | – | ↓ | ✓ | – | ↓ |
| Establishment of protein localization to mitochondrion | ✓ | – | ↓ | ✓ | – | ↓ |
| Release of cytochrome c from mitochondria | ✓ | – | ↓ | ✓ | – | ↓ |
| Protein import into mitochondrial matrix | ✓ | – | ↓ | ✓ | – | ↓ |
| Apoptotic mitochondrial changes | ✓ | – | ↓ | ✓ | – | ↓ |
| Regulation of mitochondrion organization | ✓ | – | ↓ | ✓ | – | ↓ |
| Mitochondrial translation | ✓ | – | ↓ | ✓ | – | ↓ |
| Mitochondrial gene expression | – | – | – | ✓ | – | ↓ |

Up-INT DEGs, upregulated interaction DEGs; Down-INT DEGs, downregulated interaction DEGs. ↑ indicates significant enrichment among Up-INT DEGs, ↓ indicates significant enrichment among Down-INT DEGs, ↑↓ indicates significant enrichment among both Up-INT and Down-INT DEGs, ✓ indicates significant enrichment, and – indicates no significant enrichment. DEGs, differentially expressed genes; GO, Gene Ontology; LPS, lipopolysaccharide; TNF, tumor necrosis factor.

Notably, mitochondrial pathways were broadly represented among Down-INT genes in both populations, including those related to mitochondrial organization, translation, protein import, and mitochondrial stress and apoptosis. These changes were particularly prominent in monocyte-like macrophages, which additionally showed enrichment of oxidative phosphorylation and mitochondrial gene-expression pathways among Down-INT genes. Monocyte-like macrophages also showed enrichment of hematopoiesis-regulatory pathways among Up-INT genes. Collectively, these findings reveal both shared and cell type-specific age-dependent remodeling of inflammatory, migratory, effector, and mitochondrial programs within the cardiac myeloid compartment.

### Aging modifies mitochondrial, metabolic, and effector programs in cardiac T cells and B cells

We next examined aging-dependent remodeling of the LPS response in T cells. Similar to macrophages and monocyte-like macrophages, several mitochondrial DNA-encoded transcripts showed positive age × LPS interaction effects. In contrast, Atp5g1, a nuclear DNA-encoded component of ATP synthase, showed a negative interaction effect (Figure 6A). Eif5a, which is involved in protein synthesis and cellular stress responses, also showed a negative interaction effect. GO analysis revealed distinct functional enrichment patterns between Up-INT and Down-INT genes (Figure 6B,C). Up-INT genes were enriched in αβ T-cell activation, regulation of T-cell activation, lymphocyte differentiation, and hematopoietic regulation, whereas Down-INT genes were enriched in oxidative phosphorylation, cellular respiration, mitochondrial gene expression and translation, ribosome biogenesis, and DNA replication.

**Figure 6.**
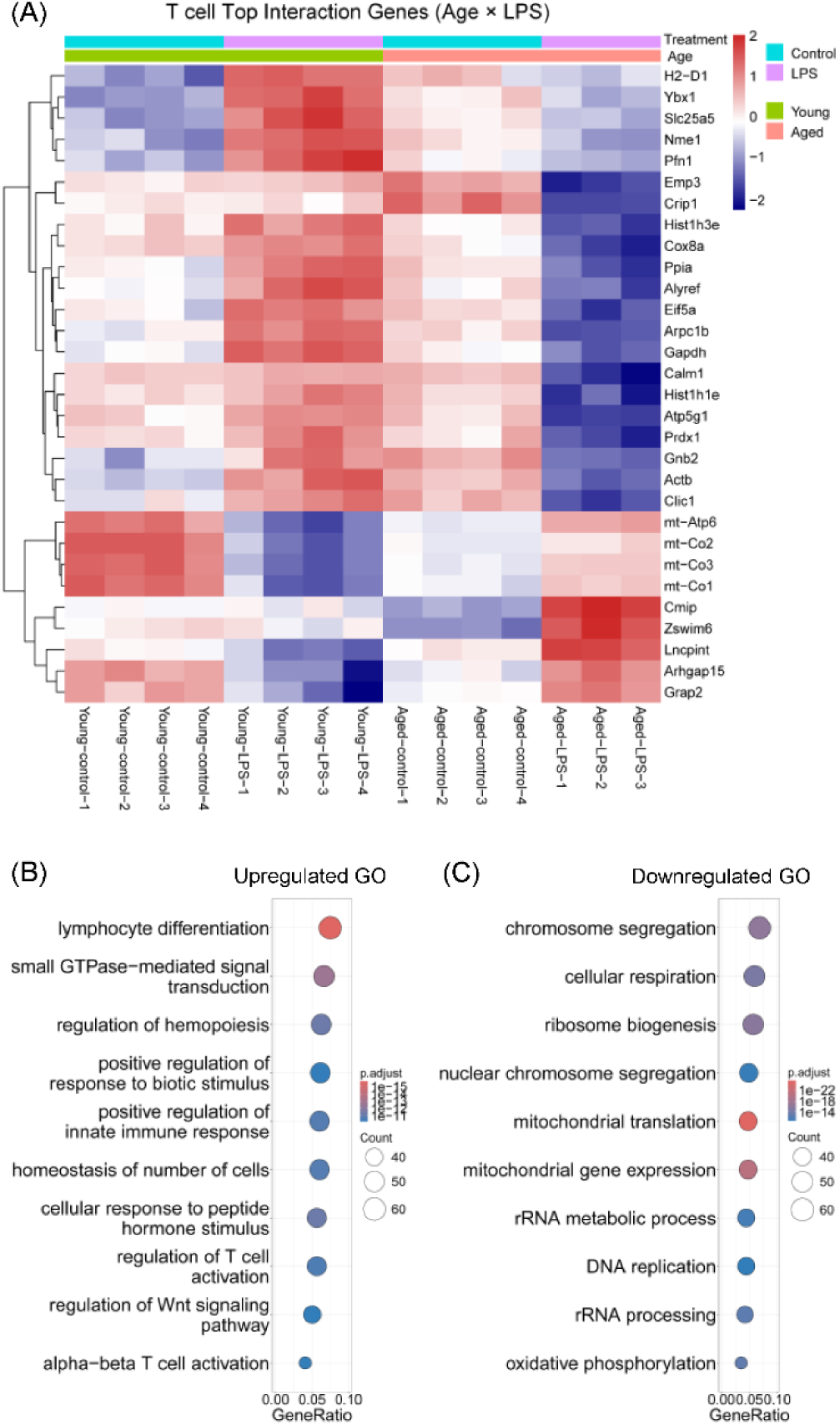
Age × LPS interaction differential expression and functional enrichment analyses in cardiac T cells. (A) Pseudobulk heatmap showing the top 30 age × LPS interaction DEGs in T cells. Genes meeting the criteria of adjusted P < 0.05 and absolute log2 fold change > 1 were ranked by adjusted P value, and the top 30 genes were selected for visualization. (B, C) Gene Ontology (GO) enrichment analysis of Up-INT genes (B) and Down-INT genes (C).

B cells exhibited a partially overlapping pattern of aging-dependent transcriptional remodeling. Several mitochondrial DNA-encoded transcripts again showed positive interaction effects, whereas Atp5f1, another nuclear DNA-encoded component of ATP synthase, showed a negative interaction effect (Figure 7A). Cd74, a component of the MHC class II antigen-presentation pathway, also showed a negative interaction effect. GO analysis demonstrated that Up-INT genes were enriched in TGF-β signaling, regulation of small GTPase-mediated signal transduction, protein phosphorylation, and actin cytoskeleton organization (Figure 7B). Down-INT genes were enriched in oxidative phosphorylation, aerobic respiration, mitochondrial gene expression and translation, respiratory-chain complex assembly, ATP metabolism, and mitochondrial organization (Figure 7C).

**Figure 7.**
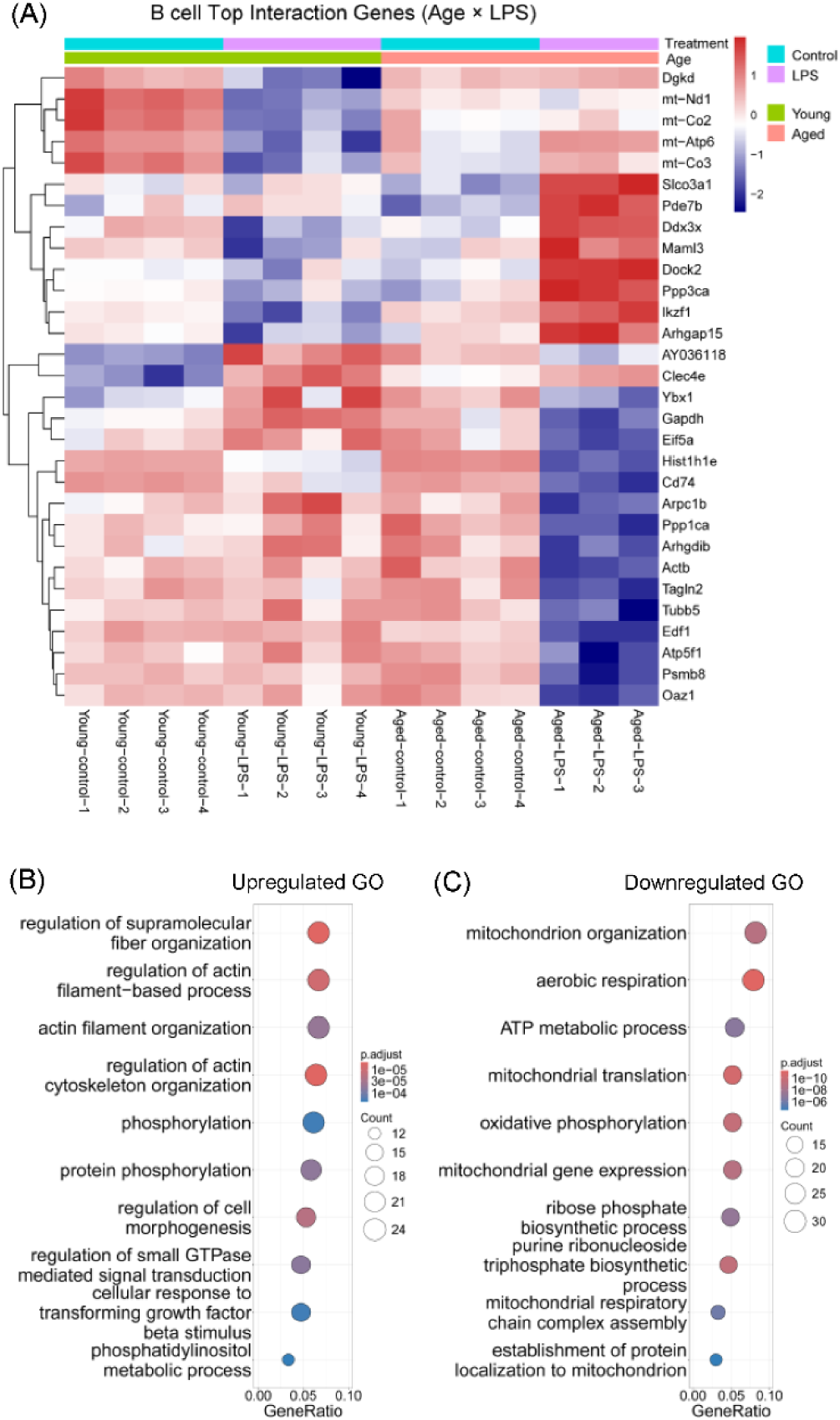
Age × LPS interaction differential expression and functional enrichment analyses in cardiac B cells. (A) Pseudobulk heatmap showing the top 30 age × LPS interaction DEGs in B cells. Genes meeting the criteria of adjusted P < 0.05 and absolute log2 fold change > 1 were ranked by adjusted P value, and the top 30 genes were selected for visualization. (B, C) Gene Ontology (GO) enrichment analysis of Up-INT genes (B) and Down-INT genes (C).

Comparison of the complete GO enrichment profiles of T cells and B cells revealed substantial overlap in mitochondrial-associated pathways (Table 2). Both populations showed Down-INT enrichment of oxidative phosphorylation, mitochondrial translation and gene expression, mitochondrial organization, respiratory-chain complex assembly, mitochondrial protein import and transport, and mitochondrial apoptotic pathways. rRNA metabolism was also enriched among Down-INT genes in both populations, whereas DNA replication was specifically enriched among Down-INT genes in T cells and ATP metabolic processes among Down-INT genes in B cells.

**Table 2.** Aging-dependent transcriptional remodeling of cardiac T cells and B cells during LPS-induced sepsis: GO enrichment of Up-INT DEGs and Down-INT DEGs.

| GO term | T cell |  |  | B cell |  |  |
|---|---|---|---|---|---|---|
|  | Down-INT | UP-INT | Summary | Down-INT | UP-INT | Summary |
| Oxidative phosphorylation | ✓ | – | ↓ | ✓ | – | ↓ |
| Mitochondrial translation | ✓ | – | ↓ | ✓ | – | ↓ |
| Mitochondrial gene expression | ✓ | – | ↓ | ✓ | – | ↓ |
| Mitochondrion organization | ✓ | – | ↓ | ✓ | – | ↓ |
| Mitochondrial respiratory chain complex assembly | ✓ | – | ↓ | ✓ | – | ↓ |
| Establishment of protein localization to mitochondrion | ✓ | – | ↓ | ✓ | – | ↓ |
| Protein import into mitochondrial matrix | ✓ | – | ↓ | ✓ | – | ↓ |
| Apoptotic mitochondrial changes | ✓ | – | ↓ | ✓ | – | ↓ |
| Mitochondrial transport | ✓ | – | ↓ | ✓ | – | ↓ |
| Regulation of release of cytochrome c from mitochondria | ✓ | – | ↓ | ✓ | – | ↓ |
| Response to oxidative stress | ✓ | ✓ | ↓↑ | ✓ | – | ↓ |
| rRNA metabolic process | ✓ | – | ↓ | ✓ | – | ↓ |
| ATP metabolic process | ✓ | ✓ | ↓↑ | ✓ | – | ↓ |
| DNA replication | ✓ | – | ↓ | – | – | – |
| Regulation of actin cytoskeleton organization | – | ✓ | ↑ | – | ✓ | ↑ |
| Actin filament organization | – | ✓ | ↑ | – | ✓ | ↑ |
| Regulation of actin filament-based process | – | ✓ | ↑ | – | ✓ | ↑ |
| Regulation of hemopoiesis | – | ✓ | ↑ | – | – | – |
| Protein phosphorylation | – | ✓ | ↑ | – | ✓ | ↑ |
| Cytotoxicity | ✓ | – | ↓ | N/A | N/A | N/A |
| T-helper 17 type immune response | – | ✓ | ↑ | N/A | N/A | N/A |
| Regulation of T-helper 17 cell differentiation | – | ✓ | ↑ | N/A | N/A | N/A |
| Antigen processing and presentation | N/A | N/A | N/A | ✓ | – | ↓ |

Up-INT DEGs, upregulated interaction DEGs; Down-INT DEGs, downregulated interaction DEGs. ↑ indicates significant enrichment among Up-INT DEGs, ↓ indicates significant enrichment among Down-INT DEGs, ↑↓ indicates significant enrichment among both Up-INT and Down-INT DEGs, ✓ indicates significant enrichment, and – indicates no significant enrichment. DEGs, differentially expressed genes; GO, Gene Ontology; LPS, lipopolysaccharide; N/A, not applicable.

In contrast, both T cells and B cells showed Up-INT enrichment of pathways related to actin cytoskeleton organization, actin filament organization, regulation of actin filament-based processes, and protein phosphorylation, indicating aging-dependent remodeling of cytoskeletal and associated signaling programs.

Beyond these shared metabolic and cytoskeletal changes, T cells showed remodeling of effector-associated pathways, with cytotoxicity-related pathways enriched among Down-INT genes and Th17-associated responses enriched among Up-INT genes. B cells, in contrast, showed Down-INT enrichment of pathways related to antigen processing and presentation.

### Representative genes illustrate major aging-dependent transcriptional patterns

To further illustrate the principal transcriptional patterns identified by the age × LPS interaction analysis, representative genes were examined across macrophages, monocyte-like macrophages, T cells, and B cells (Figure 8).

**Figure 8.**
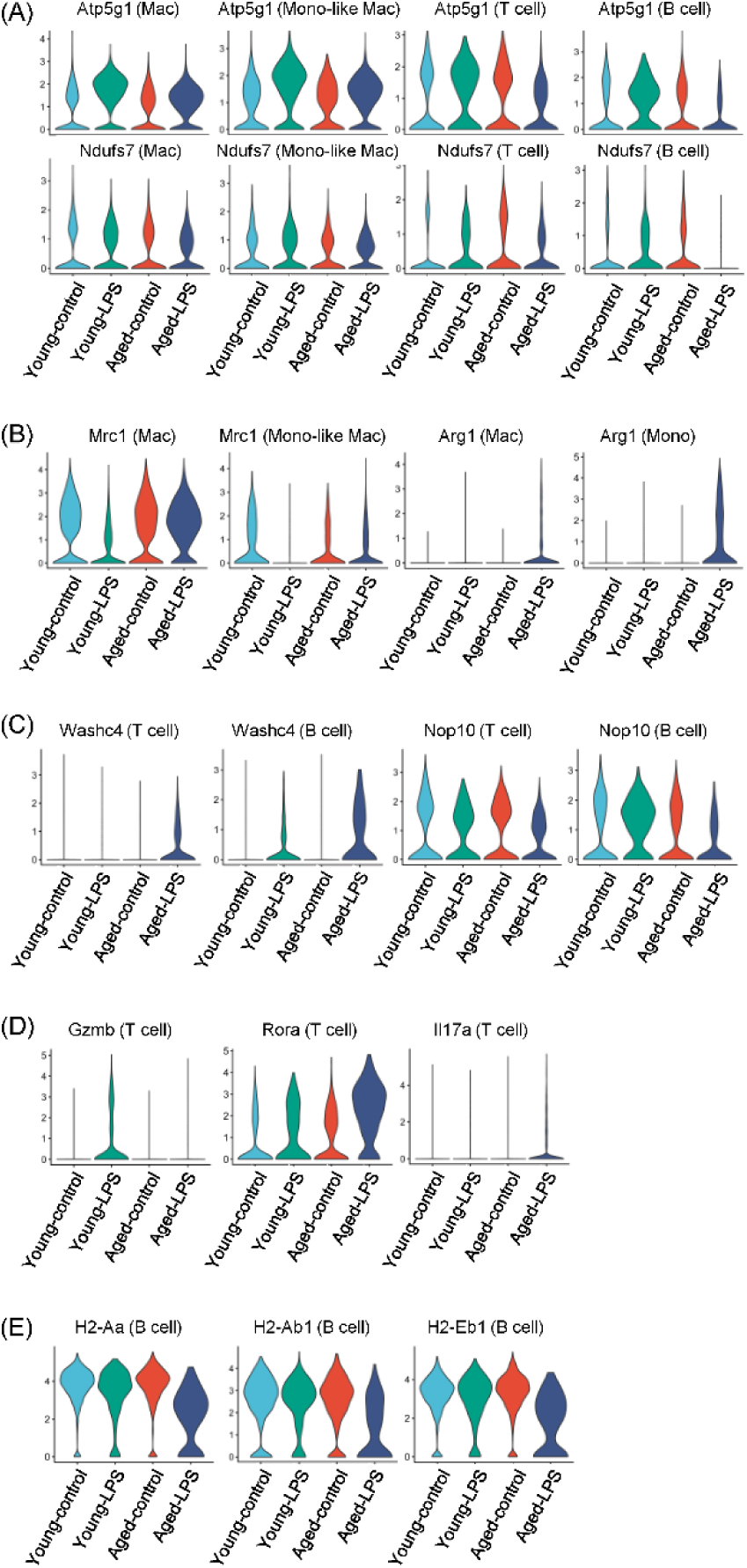
Representative age × LPS interaction DEGs illustrating major transcriptional programs remodeled by aging. Violin plots show expression of representative interaction-associated differentially expressed genes across the experimental groups. (A) Mitochondrial-associated genes Atp5g1 and Ndufs7 across macrophages, monocyte-like macrophages, T cells, and B cells. (B) Tissue-remodeling-associated genes Mrc1 and Arg1 in macrophages and monocyte-like macrophages. (C) Cytoskeletal and ribosomal RNA-processing-associated genes Washc4 and Nop10 in T and B cells. (D) T-cell effector-associated genes Gzmb, Rora, and Il17a. (E) B-cell antigen-presentation-associated genes H2-Aa, H2-Ab1, and H2-Eb1. Mac, macrophage; Mono-like Mac, monocyte-like macrophage.

The nuclear DNA-encoded mitochondrial genes Atp5g1 and Ndufs7 showed negative interaction effects across all four immune populations (Figure 8A), consistent with the broader representation of mitochondrial-associated pathways among Down-INT genes. Among myeloid populations, Mrc1 and Arg1, both associated with tissue-reparative macrophage states, showed positive interaction effects, with Mrc1 more prominent in macrophages and Arg1 in monocyte-like macrophages (Figure 8B).

In lymphocytes, Washc4, which is involved in actin organization, showed a positive interaction effect, whereas Nop10, which participates in rRNA processing, showed a negative interaction effect (Figure 8C). These patterns were consistent with the enrichment of cytoskeletal pathways among Up-INT genes and rRNA-associated pathways among Down-INT genes. T-cell effector-associated genes also exhibited age-dependent remodeling, with Gzmb, a marker of cytotoxic effector activity, showing a negative interaction effect, whereas Rora and Il17a, associated with Th17 differentiation and function, showed positive interaction effects (Figure 8D). In B cells, the antigen-presentation-associated genes H2-Aa, H2-Ab1, and H2-Eb1 showed negative interaction effects (Figure 8E), consistent with the enrichment of antigen-processing and presentation pathways among Down-INT genes.

Overall, these representative genes reinforce the major patterns identified by pathway-level analysis, including age-dependent remodeling of mitochondrial, immune-effector, tissue-remodeling, cytoskeletal, and antigen-presentation programs following LPS challenge.

### Cell–cell communication patterns among cardiac immune cell populations across experimental groups

We next examined potential interactions among cardiac immune cell populations and their possible contributions to the cardiac immune response during systemic inflammation. As shown in Figure 9A, interaction-strength circle plots revealed differences in inferred cell–cell communication patterns across the four experimental groups. At baseline, macrophages exhibited relatively strong inferred interactions in both control groups. Notably, interactions between B cells and dendritic cells or T cells were not detected in Young-control mice, whereas weak interactions were observed in Aged-control mice. Following LPS treatment, monocyte-like macrophages showed increased interaction strength in both age groups, particularly in Aged-LPS mice. Neutrophils also exhibited increased interaction strength in both LPS-treated groups relative to their respective controls, with the strongest interactions observed in Aged-LPS mice. These patterns suggested greater involvement of monocyte-like macrophages and neutrophils in the inferred communication networks following LPS challenge, particularly in aged mice. In contrast, B cells showed reduced interaction strength following LPS treatment in both age groups. Dendritic cells were absent or present at very low abundance in Aged-LPS mice, and no interactions involving dendritic cells were detected in this group.

**Figure 9.**
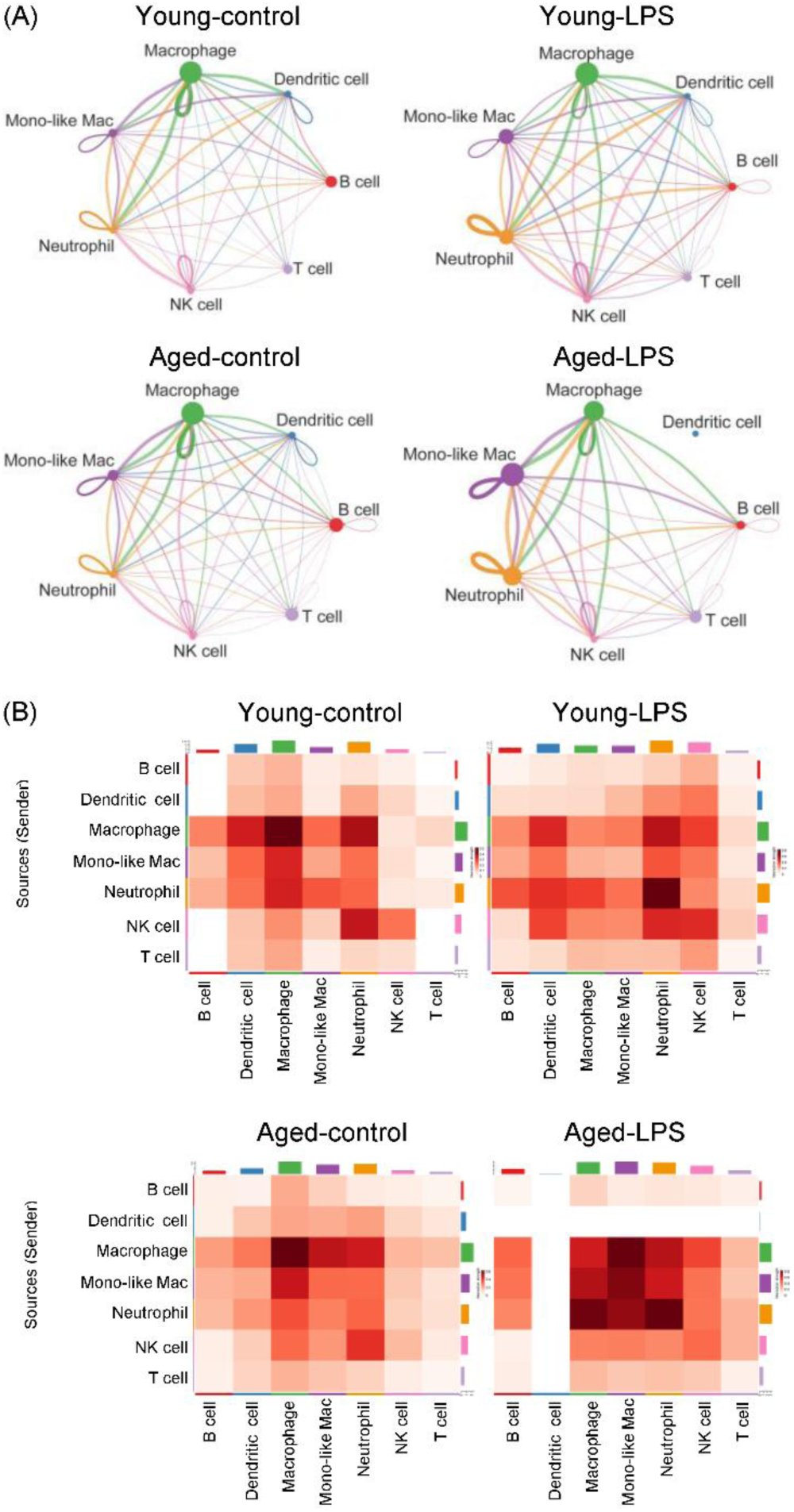
Cell–cell communication networks among cardiac immune cell populations across experimental groups. (A) Circle plots showing the strength of inferred cell–cell communication among immune cell populations in the Young-control, Young-LPS, Aged-control, and Aged-LPS groups. Nodes represent cell populations, and edges represent inferred interactions, with edge thickness proportional to interaction strength. (B) Heatmaps showing the strength of inferred interactions between immune cell populations in each experimental group. Cell–cell communication was inferred using CellChat based on normalized single-cell RNA-sequencing expression data and the mouse ligand–receptor interaction database.

Communication-strength heatmaps further revealed distinct patterns of inferred signaling among immune cell populations (Figure 9B). In both control groups, macrophages appeared to be prominent senders and receivers of inferred signals. Despite their relatively low abundance, neutrophils and NK cells showed weaker interaction strengths in Aged-control mice than in Young-control mice. In Young-LPS mice, neutrophils exhibited the strongest outgoing and incoming interaction strengths. In Aged-LPS mice, neutrophils showed strong outgoing interactions with macrophages, monocyte-like macrophages, and other neutrophils, whereas monocyte-like macrophages appeared to be the most prominent receivers of incoming signals. The absence of dendritic cell-associated interactions in the Aged-LPS heatmap was consistent with their absence or very low abundance in the corresponding circle plot (Figure 9A) and the cell-type composition shown in Figure 2C.

Overall, the inferred communication networks suggested that LPS treatment was associated with changes in interaction strength among cardiac immune cell populations, with prominent involvement of neutrophils and monocyte-like macrophages, particularly in aged mice. These findings highlight aging-associated differences in inferred cardiac immune cell communication following LPS challenge.

## DISCUSSION

### Aging drives bidirectional immune reprogramming rather than uniform immunosuppression in LPS-induced endotoxemia

In this study, we used single-cell RNA sequencing of perfused cardiac tissue to define how aging reshapes the extravascular immune response to systemic inflammation. By comparing young and aged mice under control and LPS-treated conditions and applying an age × LPS interaction model, we identified transcriptional interaction DEGs and biological programs whose responses to endotoxemia were specifically modified by aging. Notably, all four major immune populations showed substantial numbers of both positive and negative interaction DEGs, indicating that aging does not simply suppress the cardiac inflammatory response, as might be predicted by a conventional model of immunosenescence [34]. Rather, aging reshaped the response through bidirectional transcriptional changes across the cardiac immune compartment. Consistent with this pattern, GO enrichment analysis identified distinct biological programs associated with both Up-INT and Down-INT genes. Together, these findings support a model of bidirectional immune reprogramming in which aging alters the cardiac immune response to systemic inflammation rather than uniformly suppressing immune function.

### Aging reshapes cardiac immune cell composition and transcriptional states during systemic inflammation

The cardiac immune compartment underwent substantial compositional changes following LPS administration (Figure 2C). In control hearts, neutrophils were scarce, whereas LPS-induced systemic inflammation resulted in a marked influx of neutrophils into the heart, consistent with their well-established role in acute inflammation. Notably, aging induced only modest transcriptional remodeling in cardiac neutrophils (Figure 3A), suggesting that, in the LPS-stressed aged heart, the response of neutrophils was driven predominantly by changes in abundance rather than by extensive changes in their transcriptional state.

At baseline, most cardiac macrophages were resident macrophages. Aged hearts contained a smaller proportion of cardiac resident macrophages than young hearts, and NK cells were likewise reduced in proportion with aging. Together, these findings suggest that the steady-state innate immune landscape of the aged heart was characterized by a relative reduction in several immune populations compared with that of the young heart. Following LPS administration, both cardiac resident macrophages and dendritic cells decreased in proportion, consistent with previous reports that resident macrophage and dendritic cell populations can be depleted or redistributed during acute inflammation [14, 35–37]. In contrast, monocyte-like macrophages increased in proportion (Figure 2C). These cells predominantly represent monocyte-derived macrophages, and their expansion is consistent with the recruitment of circulating monocytes to the heart during LPS-induced systemic inflammation. Similarly, the increased proportion of neutrophils likely reflects their recruitment from the peripheral circulation.

The distinct responses of these immune populations to LPS can be further understood by integrating changes in relative abundance with transcriptional remodeling. Monocyte-like macrophages showed both increased relative abundance and substantial transcriptional remodeling, suggesting that these cells not only accumulated within the inflamed heart but also underwent pronounced changes in cell state. In contrast, cardiac resident macrophages decreased in relative abundance while exhibiting extensive transcriptional remodeling, indicating substantial adaptation of the remaining resident macrophage population to the inflammatory environment. Neutrophils displayed a different pattern, with a marked increase in relative abundance but relatively limited transcriptional remodeling. Thus, the cardiac immune response to systemic inflammation involved aging-dependent changes in cellular abundance and transcriptional state that varied across immune populations.

### Mitochondrial-associated transcriptional remodeling is a common feature of aging during endotoxemia

One of the most consistent findings across the four major immune populations was the enrichment of mitochondrial-associated pathways among Down-INT genes. In macrophages and monocyte-like macrophages, these pathways included mitochondrial organization, translation, protein localization and import, and mitochondrial stress- and apoptosis-related processes. T cells and B cells exhibited an even broader mitochondrial signature, encompassing oxidative phosphorylation, mitochondrial gene expression and translation, respiratory-chain complex assembly, mitochondrial transport, ATP metabolism, and related processes. The concordance of these transcriptional changes across both myeloid and lymphoid populations suggests that mitochondrial biology represents a common axis of aging-associated remodeling in the cardiac immune compartment during systemic inflammatory stress. The broad reduction in mitochondrial-associated transcriptional programs observed in aged immune cells following LPS challenge therefore raises the possibility that aging alters the capacity of cardiac immune cells to maintain or appropriately adapt mitochondrial homeostasis during systemic inflammation.

An additional finding was the increased representation of several mitochondrial DNA-encoded transcripts among Up-INT genes across the myeloid and lymphoid lineages. At first glance, this observation may appear discordant with the broader reduction in mitochondrial-associated pathways. However, increased mitochondrial DNA-encoded transcript abundance does not necessarily indicate enhanced mitochondrial function. Notably, several nuclear DNA-encoded mitochondrial genes, including Atp5g1 and Ndufs7 (Figure 8), were downregulated by aging in response to LPS across multiple cell types. Thus, the increased abundance of selected mitochondrial DNA-encoded transcripts occurred in the context of broader reduction of nuclear-DNA encoded mitochondrial programs may reflect a compensatory or stress-adaptive response in which mitochondrial-DNA encoded gene transcription was altered in an attempt to maintain components of the respiratory machinery despite disruption of broader mitochondrial homeostasis. The mitochondrial DNA-encoded response was not uniform across the experimental groups and was particularly evident in young-control and aged-LPS macrophages. Given the stringent quality-control procedures applied in this study, these transcripts are unlikely to simply reflect inadequate quality control.

The potential relevance of this finding may extend beyond cardiac immune cells. Systemic inflammatory states such as sepsis are well established to disrupt tissue oxygen and nutrient availability, imposing substantial metabolic stress on highly energy-demanding cardiomyocytes and leading to mitochondrial remodeling and dysfunction [38–41]. Our observation that cardiac extravascular immune cells similarly exhibited broad alterations in mitochondrial-associated transcriptional programs raises the possibility that mitochondrial remodeling may represent a shared feature of both cardiac leukocytes and cardiomyocytes within the inflamed myocardium.

### Aging remodels inflammatory and effector programs in cardiac macrophages

Both cardiac macrophages and monocyte-like macrophages showed enrichment of genes associated with TNF production and type I interferon responses among Down-INT genes, indicating that aging attenuated components of the inflammatory and innate immune transcriptional response to endotoxemia. In contrast, both populations showed enrichment of tissue-remodeling programs among Up-INT genes, with increased Mrc1 and Arg1 expression further supporting a reparative phenotype. Together, these findings suggest that aging shifts cardiac macrophages away from a predominantly pro-inflammatory transcriptional state toward programs associated with inflammation resolution, tissue remodeling, and repair. In the context of severe systemic inflammation, this altered balance may result in reduced inflammatory activation accompanied by greater engagement of reparative programs, potentially reshaping the macrophage response to endotoxemia in the aged heart.

### LPS-induced endotoxemia produces early adaptive immune-cell remodeling

Although LPS primarily activates innate immune pathways through the TLR4/CD14/MD-2 receptor complex, our data demonstrate substantial aging-dependent transcriptional remodeling of cardiac T and B cells 24 hours after LPS exposure. The observed changes in T-cell cytotoxicity and Th17-associated programs, together with altered B-cell antigen-processing and presentation, suggest that adaptive immune-cell programs are already being reshaped during early systemic inflammation, consistent with clinical and experimental evidence that adaptive immune responses are altered during the early phase of human sepsis and endotoxemia [42–46]. Importantly, however, most clinical studies of adaptive immune responses in sepsis have focused on circulating immune cells, whereas our analysis characterizes immune cells within the cardiac extravascular compartment.

### Aging-associated changes in cardiac immune cell communication suggest potential features of inflammaging

The cell–cell communication analysis revealed patterns that may reflect changes in the cardiac immune microenvironment across experimental groups. In both control groups, macrophages were prominent in the circle plots and appeared to be major senders and receivers in the communication heatmaps (Figure 9), consistent with the established roles of resident cardiac macrophages in immune surveillance and tissue homeostasis under steady-state conditions. Following LPS challenge, the increased representation of neutrophils and monocyte-like macrophages in the cardiac immune cell compartment (Figure 2C) was accompanied by more prominent inferred interactions involving these populations (Figure 9), particularly in aged mice. Together, these observations suggest that LPS challenge was associated with remodeling of inferred cardiac immune communication networks, with neutrophils and monocyte-like macrophages showing prominent involvement in the Aged-LPS group.

In the context of inflammaging, the comparison between Young-control and Aged-control mice revealed exploratory patterns of aging-associated differences in the cardiac immune microenvironment at baseline. Weak interactions involving B cells and dendritic cells or T cells were observed in Aged-control mice but were not detected in Young-control mice. In addition, the heatmaps suggested weaker interactions involving neutrophils and NK cells in Aged-control mice. These patterns may reflect aging-associated remodeling of cardiac immune communication at baseline, potentially involving altered lymphocyte interactions and innate immune signaling. However, because the CellChat analysis was descriptive and did not include formal statistical comparisons between groups, these observations should be interpreted as exploratory rather than as evidence of statistically significant differences. Further studies incorporating biological replicate-level communication analyses and experimental validation will be needed to determine whether these patterns represent reproducible features of cardiac inflammaging.

### Study design and scope

An important strength of this study is its prospective and well-controlled 2 × 2 factorial design incorporating age and inflammatory challenge, with biological replicates in each experimental group. This design enabled direct assessment of age × LPS interactions and allowed us to distinguish aging-dependent modification of the inflammatory response from baseline aging-related differences. Importantly, we generated an independent experimental dataset specifically designed to comprehensively characterize aging-dependent transcriptional remodeling across multiple cardiac immune-cell populations at single-cell resolution. Thus, the present study provides an experimentally grounded, cell-resolved characterization of how aging modifies the cardiac immune response to systemic inflammation. Although scRNA-seq primarily captures cellular transcriptional states rather than protein abundance or cellular function, comprehensive protein-level and functional validation of the identified transcriptional programs was beyond the scope of this study.

### Limitations

Several limitations should be considered when interpreting these findings. First, we used an LPS-induced endotoxemia model because it provides a well-controlled and reproducible inflammatory stimulus. The relatively low variability among intra-group replicates, as reflected in the heatmaps, indicates overall consistency of the transcriptional responses within experimental groups. Nevertheless, LPS-induced endotoxemia represents a model of acute systemic inflammation and does not fully recapitulate the complexity of infectious sepsis. Second, our analysis was limited to a single time point, 24 hours after LPS exposure. Longitudinal analyses across multiple stages of endotoxemia will be important for defining the temporal evolution of these aging-dependent immune responses and determining whether the observed transcriptional changes are transient or sustained. Third, neutrophils, NK cells, and dendritic cells were not included in the primary DEG analyses because these populations showed relatively few differentially expressed genes and were sparsely represented in the cardiac immune-cell compartment in at least two experimental groups. Their relatively low abundance and limited differential transcriptional responses should be considered when interpreting the overall cellular landscape of the aging-dependent cardiac immune response to endotoxemia.

## DATA AVAILABILITY STATEMENT

The datasets generated and/or analyzed during the current study will be deposited in the Gene Expression Omnibus (GEO) repository and will be made publicly available upon publication.

## ETHICS STATEMENT

All animal procedures were approved by the Institutional Animal Care and Use Committee (IACUC) of Indiana University and were conducted in accordance with institutional and national guidelines for the care and use of laboratory animals.

## AUTHOR CONTRIBUTIONS

Q.Y.: Conceptualization, Methodology, Investigation, Data curation, Formal analysis, Visualization, Writing – original draft.

H.G.: Data curation, Formal analysis.

F.F.: Methodology, Investigation.

J.D.: Investigation.

M.W.: Conceptualization, Supervision, Project administration.

## ACKNOWLEDGMENTS

The authors appreciate support of Maks Luthra on facilitating the scRNA-seq consultation with IU Center for Medical Genomics.

